# Triple labeling resolves a GPCR intermediate state by 3-color single molecule FRET

**DOI:** 10.1101/2024.10.31.621373

**Authors:** Leo Bonhomme, Ecenaz Bilgen, Caroline Clerté, Jean-Philippe Pin, Philippe Rondard, Emmanuel Margeat, Don C. Lamb, Robert B. Quast

## Abstract

The correlation of individual conformational changes in dynamic protein complexes remains challenging as most structural methods rely on averaged information over large numbers molecules. Single molecule FRET is a powerful tool for monitoring such conformational changes. When performed using three distinct probes, it enables the correlation of domain movements by providing up to three simultaneous distance measurements with high temporal resolution. Nevertheless, a major challenge lies in the site-specific attachment of three probes to unique positions within the target protein. Here, we propose an orthogonal triple-labeling strategy that is not compromised by native, reactive amino acid functionalities. It combines genetic code expansion and biorthogonal labeling of two different non-canonical amino-acids with an enzymatic self-labeling SNAP tag. We demonstrate its application by establishment of a 3-color sensor on the human metabotropic glutamate receptor 2, a dimeric, multidomain G protein-coupled neuroreceptor, and describe a previously unknown conformational intermediate state using 3-color single molecule FRET.

---

Multidomain protein complexes experience concerted conformational changes to exert their specific functions. However, a quantitative assessment of the visited conformational states and transition timescales remains challenging. Förster resonance energy transfer (FRET) is a powerful tool to derive distance information on the molecular scale from single molecules and allows one to capture transient intermediates that may be obscured by ensemble averaging (Lerner et al. 2021). Single molecule FRET (smFRET) has to date found recognizable applications for distance measurements between two probes attached to proteins (Agam et al. 2023). However, it is not possible to extract correlated motions between individual domains from such unidimensional information. To address this, smFRET employing three spectrally-separated fluorophores, can be used to simultaneously monitor and correlate three distances on a single protein or complex. This opens the possibility to identify and quantify transient intermediates in dynamic protein systems (Hohng, Joo, and Ha 2004). However, the major challenge in 3-color smFRET remains the efficient and specific conjugation of three fluorophores to selected positions on the target protein. Indeed, stochastic labelling of multiple positions using the same chemistry can complicate the interpretation of the FRET histograms due to the appearance of several FRET populations between undefined labeling positions, and thus impairs correct assignment of measured FRET efficiencies to the distance changes of interest. Previous solutions to this employed cysteine-maleimide chemistry in combination with other labeling approaches. For instance, Voss et al. combined labeling of an exposed cysteine in a protein N-terminally fused to a fluorescent protein with C-terminal oxime ligation after conversion of an intein tag to an oxyamine protein derivative (Voss et al. 2014). As an alternative, Yoo et al. first conjugated dyes to a cysteine and a site-specifically incorporated non-canonical amino acid (ncAA) and then introduced an additional, reactive cysteine through C-terminal, sortase-mediated ligation (Yoo et al. 2018). While these approaches enable site-specific triple labeling, both rely on additional C-terminal transformations before the third dye can be attached. Furthermore, fluorescent proteins are today rarely employed in quantitative smFRET measurements due to their low brightness and photo-stability. Stochastic labeling of two cysteines combined with an ncAA has also been employed for 3-color smFRET but accurate correction for the contributions of the additional FRET populations can be complicated and requires careful characterization of photo-physical properties of each of the stochastically attached dyes for either position on the target protein (Barth, Voith Von Voithenberg, and Lamb 2019; Milles et al. 2012; Voith von Voithenberg et al. 2021). Nevertheless, specificity can be promoted while using two cysteines in combination with an ncAA by tuning cysteine accessibility through ligand-binding (Agam, Barth, and Lamb 2024) or by taking advantage of differing cysteine reaction kinetics (Benke et al. 2021). While the above-mentioned approaches ingenuously tackle the challenge for their respective systems, there is still a need for more generalizable strategies that can be applied to a broader spectrum of protein targets and where the invasiveness of the labelling procedures are minimized.

In particular, the functional importance of native cysteines in many proteins, strongly limits their use to the labelling of cysteine-less proteins or requires extensive mutagenesis to remove them (Toseland 2013), and is sometimes simply incompatible with maintaining the function of the protein. An attractive alternative to cysteines are ncAAs, which barely exceed the size of proteinogenic amino acids, and allow to equip proteins with chemical handles that can subsequently be used to conjugate desired organic dyes in a site-specific and biorthogonal manner (Lee, Kang, and Park 2019). Their selective incorporation at desired positions within the protein-of-interest can be achieved through the suppression of premature stop codons, that is easily obtained by site-directed mutagenesis of the target gene. Incorporation *in cellulo* is mediated by evolved aminoacyl-tRNA synthetase/tRNA pairs that behave orthogonal to the host protein synthesis machinery.

Here, we present a site-specific triple labelling strategy based on two different ncAAs and a SNAP tag and applied it to uncover an intermediate state during activation of a G protein-coupled receptor (GPCR, **Fig. 1**). We focused on the human metabotropic glutamate receptor 2 (mGlu2), a class C GPCR essential in the regulation of neuronal excitability and synaptic transmission (Niswender and Conn 2010). Cryo-electron microscopy structures of this dimeric receptor (mediated by a disulfide bridge between the two protomers, **Fig. 1**) point at a transition from an inactive, resting open (R_O_) state to an active closed state (A_C_) during their activation (Du et al. 2021; Lin et al. 2021; Seven et al. 2021). This involves a ligand-induced closure of the Venus flytrap (VFT) domains (open o ↔ c closed transition), a reorientation of the two subunits relative to each other bringing the lower lobes of the VFT domains in closer proximity (resting R ↔ A active transition) and changes to the interface of adjacent 7 transmembrane (7TM) domains (**Fig. 1**). Using a 2-color smFRET sensor of the o ↔ c transition, we recently revealed that saturating concentrations of the natural full agonist glutamate lead to an efficient closure of the VFT domains (Lecat-Guillet et al. 2023). However, by monitoring the R ↔ A transition, we discovered the co-existence of at least two states corresponding to the resting and active orientations. These data suggest an agonist induced dynamic equilibrium between three states including an intermediate state between the R_O_ and the A_C_ state (**Fig. 1**). However, a direct and simultaneous correlation of VFT domain closure and reorientation would be necessary to identify and characterize this intermediate state.

**Figure 1:**
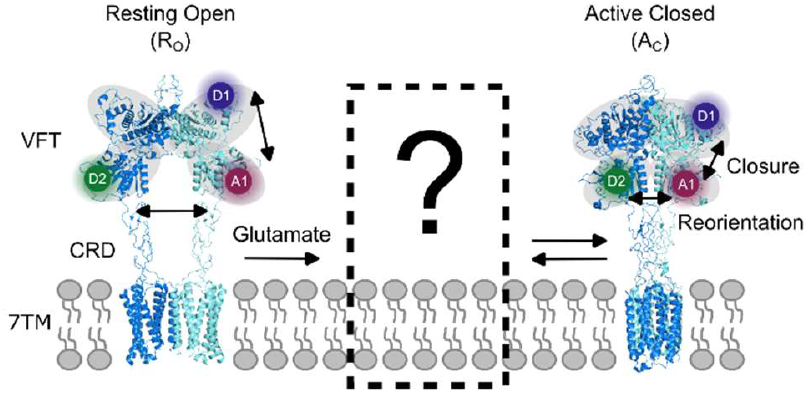
Major conformational changes during mGlu2 activation. Cryo-EM structures of (left) the resting open (R_O_) state with Venus flytrap (VFT) domains open, the lower lobes and cysteine-rich domains (CRDs) separated, and an interface mediated by transmembrane domain helices IV (PDB 7EPA) and (right) active closed (A_C_) state with the VFT domains closed, the lower lobes and CRDs in closer proximity and a slightly twisted dimer interface mediated by 7TM helices VI and VII (PDB 7EPB). We previously proposed that the glutamate-induced transition from the R_O_ to the A_C_ state might involve a resting closed (R_C_) intermediate state (Lecat-Guillet et al. 2023) but direct evidence for its existence is currently missing. The upper and lower lobes of the VFT domains are shaded in grey. The labeling positions required to establish a 3-color smFRET sensor are indicated by colored circles (Donor 1 (D1): purple; Donor 2 (D2): green; Acceptor 1 (A1): red).

To investigate potential correlations between the different domains, we developed a triple labeling strategy based on orthogonal, cotranslational incorporation of two distinct reactive ncAAs using two distinct stop codons together with fusion to a genetically encoded self-labelling SNAP tag (Keppler et al. 2003). In the first protomer of mGlu2, a SNAP-tag is fused to the N-terminal and we incorporated a p-propargyloxy-L-phenylalanine (PrF) in response to the Amber codon (TAG) at position 248 within the lower lobe (**Fig. 2A**). To the second protomer, no SNAP tag was fused and a trans-cyclooct-2-en-L-lysine (TCOK) was incorporated in response to an Ochre codon (TAA) at position 258 within the lower lobe (**Fig. 2A**). The 258 position was chosen to improve protein yields due to an inefficient incorporation of TCOK in response to TAA at the 248 position (**Supplementary Fig. 1**). PrF and TCOK were incorporated by co-expression of the receptor genes harboring the respective premature stop codons with an engineered *B. stearothermophilus* tyrosyl*-*tRNA_CUA_/*E. coli* tyrosyl-tRNA synthetase pair (PrFRS-tRNA_CUA_) (Lecat-Guillet et al. 2023) and an engineered M15 pyrolysyl-tRNA_UUA_ or -tRNA_UCA_/*M. barkeri* pyrolysyl-tRNA synthetase pair (PylRS-tRNA_UUA_) (Serfling et al. 2018) in HEK293T cells, respectively (**Fig. 2A**). To specifically label dimers with a single set of these three functionalities, we employed our previously described C1/C2 system derived from the GABA_B_ receptor quality control system (Brock et al. 2007; Lecat-Guillet et al. 2023). Subsequent labelling was performed directly on living mammalian cells using commercially available reagents for i) the O^6^-alkylguanine-DNA-alkyltransferase activity of the SNAP tag (**Fig. 2B**, D1 = Atto488), ii) strain-promoted inverse electron-demand Diels-Alder cycloaddition (SPIEDAC, **Fig. 2B**, D2 = Cy3B), and iii) copper-catalyzed azide-alkyne cycloaddition (CuAAC, **Fig. 2B**, A1 = AF647). Of note, this approach is not compromised by natively present reactive amino acid functionalities and does not require post-expression protein processing before dye conjugation.

**Figure 2:**
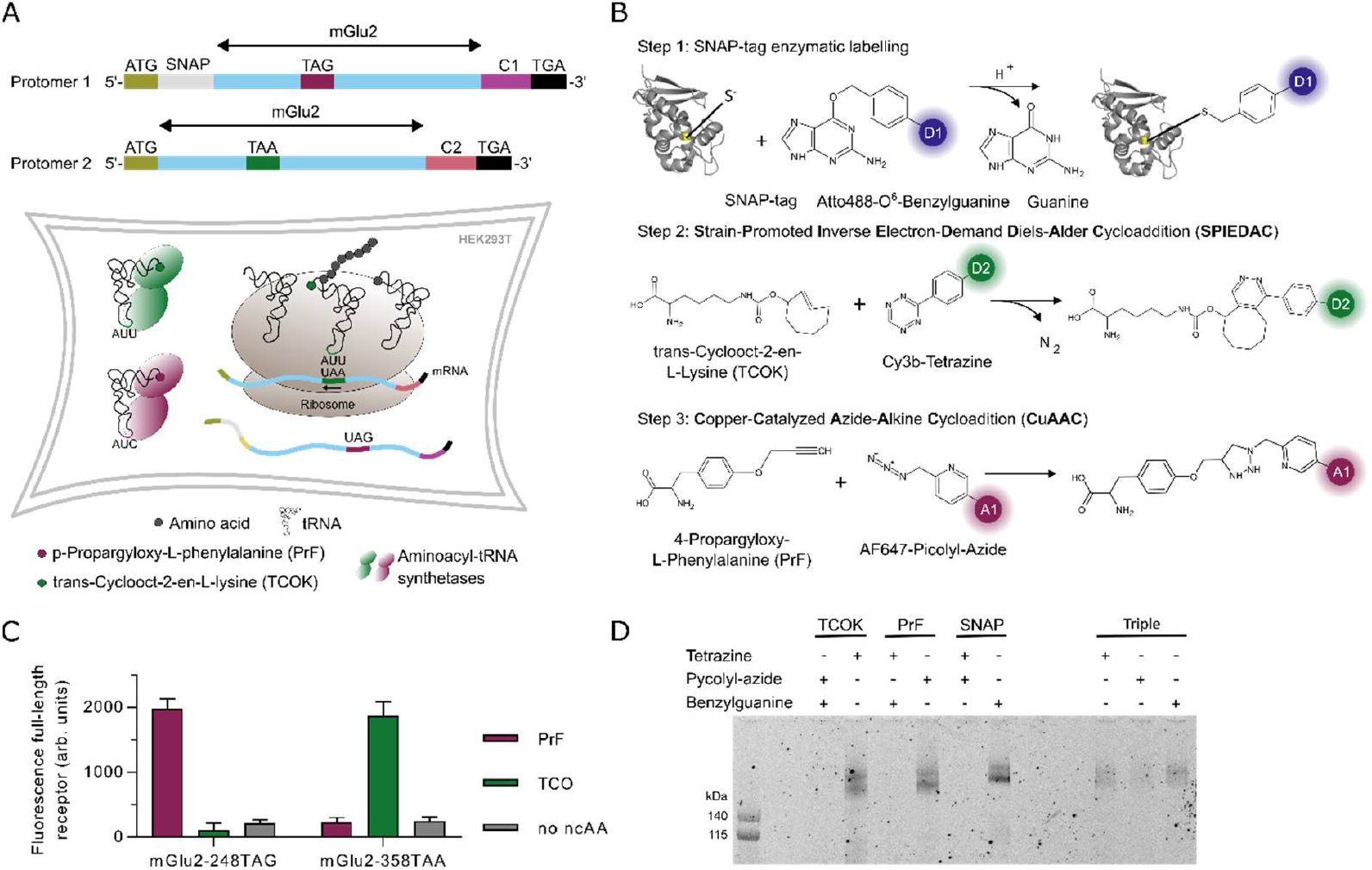
Orthogonal triple labeling strategy. (A) Top: Schematic representation of the mGlu2 protomer genes expressed to produce the 3-color smFRET sensor. Important features of the protomers are highlighted, including ATG start codons, SNAP tag, premature TAG and TAA stop codons within the mGlu2 gene, the GABA_B_ C1 and C2 tails and the final TGA stop codons. Bottom: Schematic representation of cotranslational double ncAA incorporation in HEK cells. (B) Three-step site-specific labeling reactions of mGlu2 receptors in the membrane of living cells prior to solubilization: 1) SNAP-tag labeling using Atto488-0^6^-benzylguanine; 2) TCOK labeling using Cy3B-Tetrazine; 3) PrF labeling using AF647-Picolyl-Azide. (C) Orthogonal suppression of TAG and TAA stop codons. Fluorescence signal of HEK cells expressing mGlu2 plasmid bearing either a TAG or a TAA premature stop codon and an N-terminal SNAP fusion protein, labelled at the cell surface with Lumi4-Tb-BG. Only full-length receptors expressed through successful ncAA incorporation are presented at the cell-surface and labeled. All expressions were performed with both synthetase/tRNA pairs in the absence or presence of either ncAA. (D) Evaluation of the labeling specificity. Receptors with incorporated TCOK, PrF or SNAP fusion protein were subjected to the respective labelling reactions as indicated with (+) using Cy3B-Tetrazine, AF546-Picoly-Azide and/or Cy3B-BG.

We first verified the orthogonality of the two aminoacyl-tRNA synthetase/tRNA pairs to incorporate PrF in response to TAG suppression in combination with TAA suppression for TCOK incorporation. For this we took advantage of the fact that termination at each of the premature stop codons, located within the extracellular domains, does not result in translation of the full-length receptors. Hence, only full-length receptors that comprise the 7TM domains, resulting from incorporation of ncAA, will be membrane-embedded and presented on the surface of cells. Specific labelling of cell surface-presented full-length mGlu2, through an N-terminal SNAP-tag using the membrane impermeable Lumi4-Tb-BG, served as a reporter for proper ncAA incorporation and showed the expected specificity a TAG template for PrF and a TAA template for TCOK-mediate full-length receptor expression (**Fig. 2C**).

Next, we addressed the specificity of dye conjugation to the two ncAAs and the SNAP-tag (**Fig. 2E**). Cross-reactivity leading to attachment of dyes to positions other than the desired ones would compromise the specificity of the labeling and thereby complicate assignment of observed FRET distributions to the conformational states of interest. Therefore, we expressed the two synthetase/tRNA pairs in the presence of both ncAAs together with the receptor gene harboring either: 1) only a premature TAA (**Fig. 2D**, TCOK), 2) only a premature TAG (**Fig. 2D**, PrF), 3) no premature stop codon but an N-terminal SNAP-tag (**Fig. 2D**, SNAP) or 4) both premature TAA and TAG, and the SNAP-tag (**Fig. 2D**, Triple). We then performed either the desired labelling reaction (SNAP with O^6^-Benzylguanine, TCOK with tetrazine and PrF with picolyl-azide) or a 2-step double-labeling with the two other reactions, which are not supposed to label the single functionalized samples. In-gel fluorescence demonstrated that the TAA template is required for TCOK incorporation and SPIEDAC conjugation, the TAG template is required for PrF incorporation and CuAAC conjugation and the SNAP-tag is required for the alkyltransferase self-labeling. Correspondingly, no labeling was observed for the controls highlighting the exclusive incorporation of the ncAA in response to their annotated premature stop codons, while not being incorporated in response to any sense codons. Moreover, the three reactions are highly selective leading to perfect orthogonality of our triple labeling approach. Furthermore, weak but detectable bands confirmed the successful labeling of the triple functionalized receptors for each of the three orthogonal reactions (**Fig. 2E**).

We then characterized two sensors monitoring the VFT closure (o ↔ c, **Fig. 3A**) and lower lobe reorientation (R ↔ A, **Fig. 3B**) using 2-color smFRET experiments. For each sample, two of the three positions were labeled using the same positions, strategies, and dyes as were used for the subsequent 3-color sensor. The results of the 2-color FRET experiments suggested a nearly complete closure of the VFT domains by a transition from a single low FRET (**Fig. 3A**, Apo, grey) to a medium FRET population (**Fig. 3A**, Glutamate, yellow) but only a partial stabilization of the fully reoriented active state at saturating Glutamate concentrations (**Fig. 3B**). These findings are in very good agreement with our previous observations with similar, but distinct FRET sensors, pointing to the existence of an intermediate in the presence of glutamate, being in dynamic equilibrium between resting and active orientations at timescale slower than the ∼3 ms observation time (Lecat-Guillet et al. 2023).

**Figure 3:**
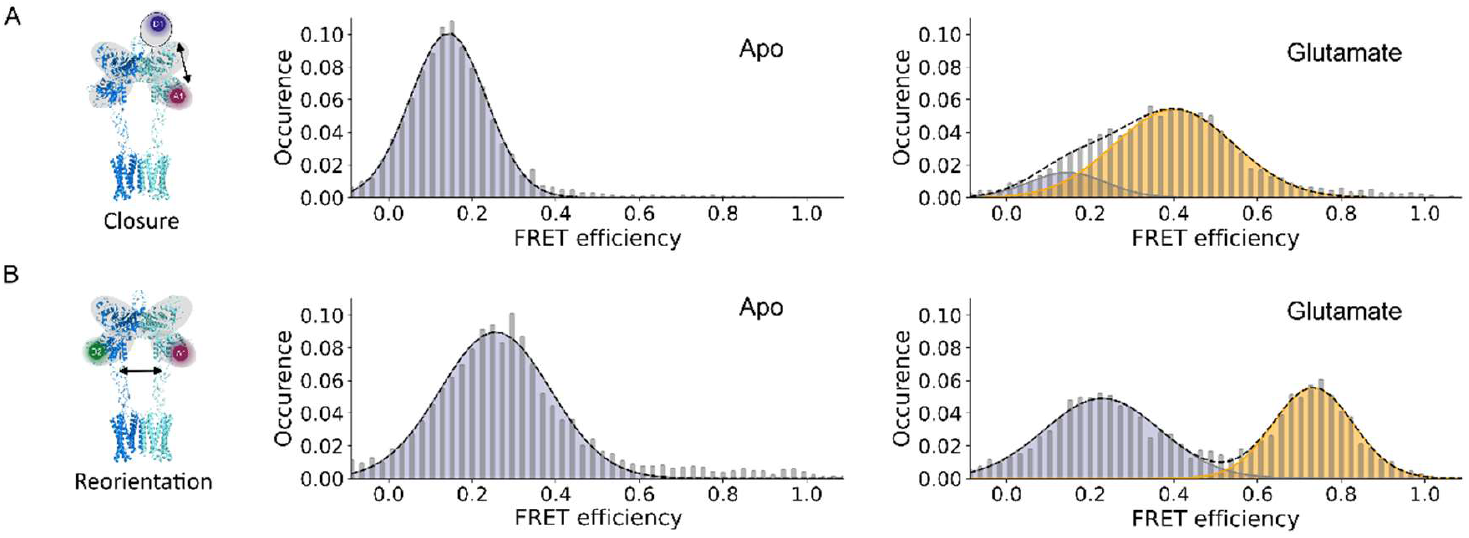
2-color smFRET experiments suggest efficient VFT domain closure but incomplete stabilization of the reoriented state. Histograms of VFT closure (A) and reorientation (B) sensors in the apo condition and in the presence of a saturating Glutamate concentration (10 mM).

Next, we measured our triple-labeled sensor using 3-color smFRET (**Fig. 4A and B**) to directly correlate VFT closure (o ↔ c, FRET efficiency BR) and reorientation (R ↔ A, FRET efficiency GR) on individual single molecules. As expected, a major low FRET state resulting from the R_O_ conformation in the absence of ligand was observed for both FRET pairs (**Fig. 4A**), when the two FRET efficiencies were displayed in a 2-dimensional plot (E_BR_ = 0.2 and E_GR_ = 0.12). In the presence of glutamate, the R ↔ A sensor revealed the two expected conformations corresponding to the resting and active orientations with similar relative population (55% vs. 45% respectively, **Fig. 4B**, FRET efficiency GR). Likewise, the o ↔ c sensor revealed two populations (**Fig. 4B**, FRET efficiency BR), with the high FRET state corresponding to the closed VFT domains in the active orientation. The 3-color smFRET data allowed us to separate these two species and determine their individual FRET efficiencies BR, reporting on the closure of the VFT domain. For the active species (high FRET efficiency GR), the FRET efficiency BR corresponded to the closed VFT domain as expected (E_BR_ = 0.46). However, to our surprise, for the species remaining in the resting state (low FRET efficiency GR), we found a FRET efficiency BR that appeared intermediate between those of the open and the closed conformation (E_BR_ = 0.34 vs. E_BR, open_= 0.2 and E_BR, closed_ = 0.46).

**Figure 4:**
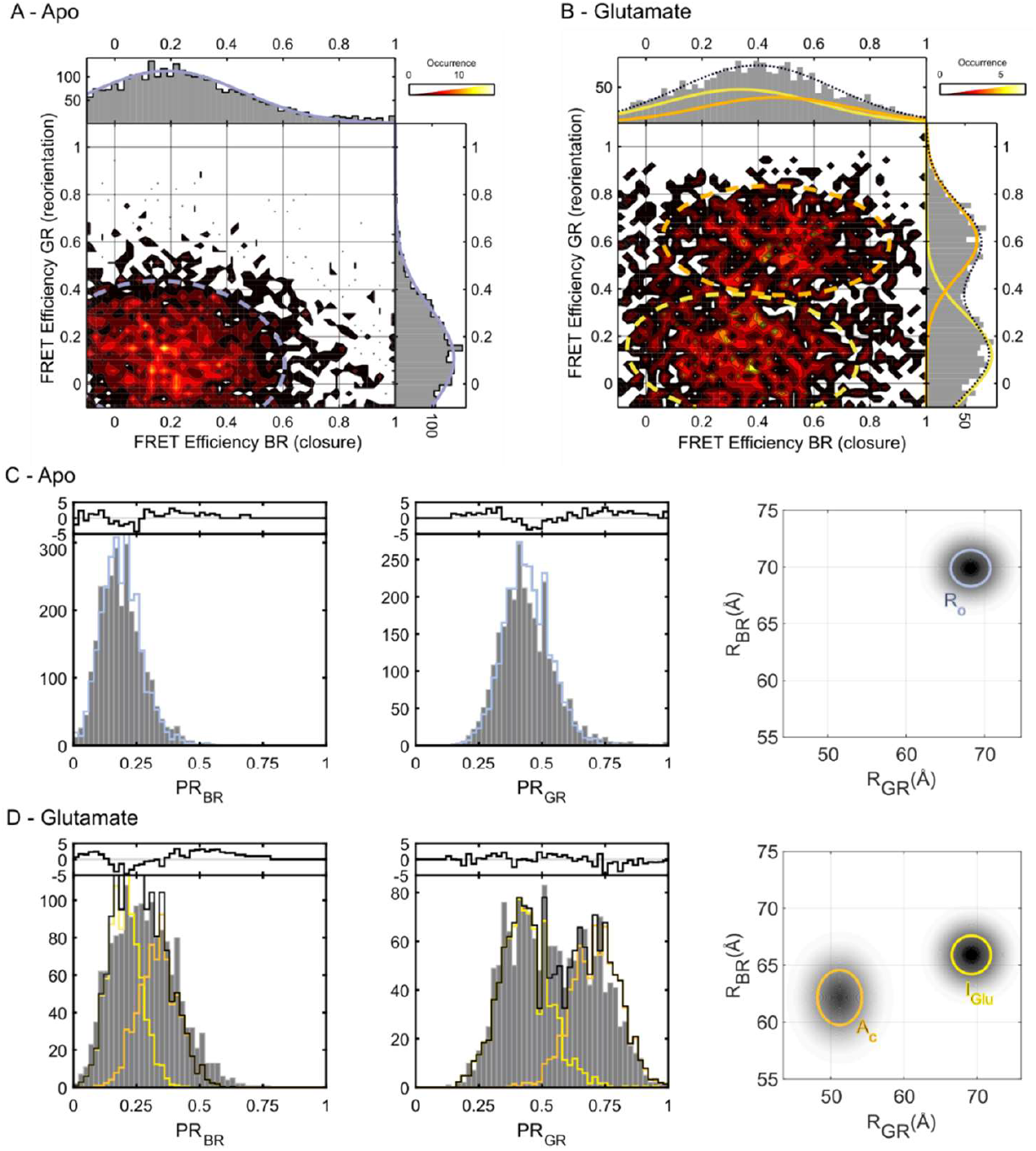
3-color smFRET experiments unveil coordinated rearrangements of the VFT domains of mGlu2 and an unknown conformational intermediate. A-B) Two-dimensional projections of VFT domain closure (FRET efficiency BR) and reorientation (FRET efficiency GR) in the absence (A) and presence of a saturating Glutamate concentration (B). C) One-dimensional projections of the proximity ratio for VFT domain closure (PR_BR_, left) and reorientation (PR_GR_, center) and two-dimensional apparent distance distribution histogram (right) extracted by PDA analysis for the apo state. (D) One-dimensional projections of the proximity ratio for VFT domain closure (PR_BR_, left) and reorientation (PR_GR_, center) and two-dimensional apparent distance distribution histogram (right) extracted by PDA analysis under the influence of a saturating glutamate concentration (10 mM). Two populations are sufficient to describe the 3-color data, which is clearly observable in the PR_GR_ projection (D, center) but more difficult to detect for the PR_BR_ data (D, left). However, the PDA analysis reveals a clear reduction in the distance between upper and lower lobes (R_BR_) between the apo (C, right) and glutamate condition (D, right) indicating a slight closure of the VFTs.

To quantitatively analyze the 3-color smFRET data, we performed our previously established 3-color photon distribution analysis (3C-PDA) using either a 1 or 2-state model to reinforce the idea that this shift corresponds to a change in the inter-dye distance distribution (Barth et al. 2019). Consistent with the 2-color data (**Fig. 3A and B**), the 1-dimensional projection of the o ↔ c sensor (PR_BR_) demonstrated the expected shift from a low FRET (**Fig. 4C**, left panel) to higher FRET values in the presence of glutamate (**Fig. 4D**, left panel), indicative of VFT closure. Accordingly, the 1-dimensional projection of the R ↔ A sensor revealed a single low FRET state (**Fig. 4C**, middle panel) that is approximately half depopulated, giving rise to a high FRET state in the presence of glutamate (**Fig. 4D**, middle panel). Thanks to the 3-color approach, we found that only two species were sufficient to describe the data in the presence of glutamate and we were able to separate out the peak in the PR_BR_ histogram into two conformations (**Fig 4D**, left panel). The population observed with higher PR_GR_ values can be assigned to the A_c_ state and has a corresponding shift in the PR_BR_ value indicative of full VFT domain closure, which corresponds to a shorter distance R_BR_ between the upper and lower lobes (**Fig. 4D**, orange population in right panel). The conformation with the lower PR_GR_ value corresponds to an increased distance between the lower lobes and has an intermediate PR_BR_ value, indicative of a slightly reduced distance R_BR_ between upper and lower lobes (**Fig. 4D**, yellow in right panel). This distance is slightly shorter than the one obtained for the R_O_ state measured under apo conditions (**Fig. 4C**, blue in right panel). This implies that the glutamate-bound VFT domains in the resting orientation adopt a slightly closed conformation, that is intermediate between the fully open state observed in the apo condition (R_O_), and the glutamate-induced fully closed state observed for the receptors in the reoriented conformation (A_C_). Insteat of establishing a dynamic equilibrium between R_C_ and A_C_ conformations, as we previously postulated (Lecat-Guillet et al. 2023), the 3-color data clearly show that glutamate-induced activation of mGlu2 receptors involves a previously unknown intermediate state (I_Glu_), which exists in equilibrium with the A_C_ state.

Two structural studies of the mGlu5 receptor recently proposed the presence of the R_CC_ (i.e. with both VFT domains closed) intermediate state (Cannone et al. n.d.; Krishna Kumar et al. 2024) we had postulated before for mGlu2 (Lecat-Guillet et al. 2023). However, based on our data presented here this state seems not to be significantly populated by mGlu2 under equilibrium conditions in a carefully selected detergent micelle environment, for which we previously demonstrated functional allosteric modulation and G protein-coupling (Cao et al. 2021). In addition, intermediates with one VFT domain in the open and the other in the closed conformation but lower lobes separated as seen for the resting state have been proposed for the mGlu2 receptor (Seven et al. 2021; Zhu et al. 2024). However, if such a conformation was significantly populated and stable under our conditions, this should be reflected by the presence of two well-defined FRET populations corresponding to resting states resulting from open-closed transitions at timescales slower than the observation time. Instead, our 3-color PDA analysis describes the data well using only two populations, representing the intermediate closed (I_Glu_) and fully closed (A_C_) states.

In summary, we demonstrated the discovery of a previously unknown intermediate state of the mGlu2 receptor when activated by its natural full agonist glutamate, which was obscured in classical 2-color smFRET experiments before. This discovery was made possible only by the 2-dimensional resolution of 3-color smFRET experiments. To perform these experiments on such a multidomain, multimeric, human neuroreceptor, it was necessary to establish a new and sophisticated orthogonal triple-labelling strategy, to overcome previous limitations related to the use of cysteine-maleimide labeling. Our strategy based on expression in mammalian cells, now opens the possibility to perform triple labeling on a broader range of proteins and complexes without being limited by natively present reactive amino acids. This opens up the possibility to study other mGlu receptor homo-but also heterodimers as well as other class C GPCRs and multimeric membrane proteins to elucidate symmetric and asymmetric structural features and provide deeper insights how conformational states and dynamics regulate protein function through concerted structural rearrangements.

## Material and method

### Chemicals

Aminoguanidine hydrochloride, copper(II) sulfate, (+)-sodium L-ascorbate, tris[(1-benzyl-1H-1,2,3-triazol-4-yl)methyl]amine, THPTA, L-glutamate, and chemicals used for synthesis were purchased from Sigma-Aldrich (St. Louis, MO, USA) unless stated otherwise. SNAP-Surface Alexa Fluor 488 was purchased from NEB (Evry, France). O-Propargyl-L-tyrosine hydrochloride (PrF) was from Iris Biotech GmbH (Marktredwitz, Germany) and trans-Cyclooct-2-en - L-Lysine (TCOK) from Sirius Fine Chemicals GmbH (Bremen, Germany). 2-(4-((bis((1-(tert-butyl)-1H-1,2,3-triazol-4-yl)methyl)amino)methyl)-1H-1,2,3-triazol-1-yl)acetic acid (BTTAA), AF647-Picolyl-Azide and AF546-Picolyl-Azide were purchased from Jena Bioscience (Jena, Germany). Cy3B tetrazine was obtained from AAT Bioquest (Pleasanton, USA). Lauryl maltose neopentyl glycol (LMNG) and cholesteryl hemisuccinate tris salt (CHS) were purchased from Anatrace (through CliniSciences, France). Glyco-diosgenin (GDN) was purchased from Avanti Polar Lipids through Merck.

### DNA constructs

The pcDNA plasmid encoding human mGlu2 with N-terminal FLAG (Sigma-Aldrich) and SNAP-tags (New England Biolabs) was a gift from Revvity (Codolet, France).

The engineered GABA_B_ C-terminal tails were obtained using restriction enzymes from the pRK5 plasmids encoding for rat mGlu2 (rat-mGluR2-C1KKXX and rat-mGluR2-C2KKXX) described in (Doumazane et al. 2011), purified by agarose electrophoresis and ligated into to human mGlu2 gene cut using the same restriction enzymes.

Premature stop codons were introduced using the QuickChange Lightning Site-directed Mutagenesis kit from Agilent Technologies (Santa Clara, CA, USA) according to the manufacturer’s protocol. Final stop codons were changed to TGA following the same protocol. N-terminal FLAG and SNAP-tags were removed from the mGlu2-258TAA-C2KKXX construct using the In-Fusion® HD cloning kit (Takara Bio Europe) according to the manufacturer’s protocol.

The pNEU-hMbPylRS(AF)-4xU6M15 plasmid used for incorporation of TCOK in response to TAA and the pIRE4-PGK-ePrFRS plasmid used for incorporation of PrF in response to TAG were a kind gift of I. Coin (Universität Leipzig, Germany).

### Protein expression, labelling and purification

All mGlu2 receptors were expressed in adherent HEK293T cells (American Type Culture Collection CRL-3216, LGC Standards S.a.r.l., France) cultured in Dulbecco’s modified Eagle’s medium (DMEM; Thermo Fisher Scientific) supplemented with 10% fetal bovine serum (FBS; Sigma-Aldrich) at 37°C with 5% CO_2_. HEK cells were transfected using JetPrime transfection reagent (Polyplus-transfection SA, Illkirch-Graffenstaden, France).

For the selective ncAA incorporation essay, HEK293T cells were cultured in 96-well F-bottom black plates (Greiner Bio-One). A total of 75,000 cells were seeded in 100 μl of DMEM per well 17 hours before transfection. Cells transfected with vectors coding for SNAP-mGlu2 with either the A248TAG or the R358TAA mutation were co-transfected with pIRE4-PGK-ePrFRS and pNEU-hMbPylRS(AF)-4xU6M15 using 250 ng of total DNA per well with a 2:1:1 ratio in 30 μl of JetPrime buffer (Polyplus). A total of 0.5 μl of JetPrime Reagent per well was added and incubated for 25 min at room temperature. Last, the transfection mixtures were added to the cells and incubated at 37°C for 6 hours. After 6h, medium was replaced with 100 μl fresh DMEM per well, supplemented with either 0.2 mM TCOK or 0.3 mM PrF. The medium was further exchanged with fresh ncAA-containing medium after 24 and 48 h. 72h after transfection SNAP-tag labelling was performed on adherent cells at 37°C and 5% CO_2_ for 2 h using a final concentration of 100 nM SNAP-Lumi4-Tb (PerkinElmer, Codolet, France) in Gibco DMEM GlutaMax without phenol red, supplemented with GlutaMAX and pyruvate (Thermo Fisher Scientific, France). Following labelling, excess dye was removed by three cycles of washing with DPBS without Ca^2+^ and Mg^2+^ (Thermo Fischer Scientific, France) at ambient temperature. Live-cell fluorescence reading was performed in 50 μl acquisition buffer (20 mM Tris-HCl pH 7.4, 118 mM NaCl, 1.2 mM KH2PO4, 1.2 mM MgSO4, 4.7 mM KCl and 1.8 mM CaCl2) on SPAK20M plate reader (Tecan) at excitation 320/25 nm and emission 620/20 nm.

For the labelling specificity assay, HEK293T cells were cultured in standard flat 6-well TC-plate (Sarstedt). A total of 1,200,000 cells were seeded in 2 ml of DMEM per well 17 hours before transfection. For the PrF condition, cells were transfected with vectors coding for mGlu2-C2KKXX, mGlu2-A248TAG-C1KKXX, pIRE4-PGK-ePrFRS and pNEU-hMbPylRS(AF)-4xU6M15 in a 1:3:4:4 ratio using 2μg of DNA per well in 200 μl of JetPrime buffer (Polyplus) per well. For the TCOK condition, cells were transfected with vectors coding for mGlu2-C1KKXX, mGlu2-L258TAA-C2KKXX, pIRE4-PGK-ePrFRS and pNEU-hMbPylRS(AF)-4xU6M15 in a 1:3:4:4 ratio using 2μg of DNA per well in 200 μl of JetPrime buffer (Polyplus) per well. For the SNAP condition, cells were transfected with vectors coding for SNAP-mGlu2-C1KKXX, mGlu2-C2KKXX, pIRE4-PGK-ePrFRS and pNEU-hMbPylRS(AF)-4xU6M15 a 1:1:1:1 ratio using 2μg of DNA per well in 200 μl of JetPrime buffer (Polyplus) per well. For the triple condition, cells were transfected with vectors coding for SNAP-mGlu2-A248TAG-C1KKXX, mGlu2-L258TAA-C2KKXX, pIRE4-PGK-ePrFRS and pNEU-hMbPylRS(AF)-4xU6M15 in a 1:1:1:1 ratio using 2μg of DNA per well in 200 μl of JetPrime buffer (Polyplus) per well. 4 μl of JetPrime Reagent per well were added to the transfection mixtures and incubated for 25 min at room temperature. Last, the transfection mixtures were added to the cells and incubated at 37°C for 6 hours. After 6h, medium was replaced with 2 ml fresh DMEM supplemented with 0.2 mM TCOK and 0.3 mM PrF. The medium was further exchanged with fresh ncAA-containing medium after 24 and 48 h. 72h after transfection SNAP-tag labelling was performed on adherent cells at 37°C and 5% CO_2_ for 2h using a final concentration of 600 nM SNAP-Cy3B, excess dye was removed by three cycles of washing with DPBS without Ca^2+^ and Mg^2+^ at RT. TCOK labelling was performed right after SNAP-tag labelling using a final concentration of 1 mM Cy3B tetrazine in acquisition buffer (20 mM Tris-HCl pH 7.4, 118 mM NaCl, 1.2 mM KH_2_PO_4_, 1.2 mM MgSO_4_, 4.7 mM KCl and 1.8 mM CaCl_2_) incubated for 15 min at 37°C. Excess dye was removed by three cycles of washing with DPBS at room temperature. PrF labelling was done last using a final concentration of 10 mM AF546-Picolyl-Azide, 1.5 mM Aminoguanidine, 1.98 mM BTTAA, 0.36 mM CuS0_4_, 2 mM Na-Ascorbate in acquisition buffer incubated for 15 min at 37°C. Excess dye was removed by three cycles of washing with DPBS. For crude membrane fractions preparation, adherent cells were detached mechanically using a cell scraper in DPBS and collected at 500 ×g and 22°C for 5 min. Subsequently, cells were re-suspended in cold hypotonic lysis buffer (10 mM HEPES pH 7.4 and cOmplete™ protease inhibitor, Roche), frozen, and stored at −80°C. After thawing, cells were passed through a 200 μl pipette tip 30 times on ice. After two rounds of centrifugation at 500 ×g and 4°C for 5 min, the supernatant was centrifuged at 21,000 ×g and 4°C for 30 min to collect crude membranes. The pellets were washed once with acquisition buffer.

Receptors were solubilized using 10 μl of acquisition buffer containing 1% LMNG (w/v) and 0.1% CHS Tris (w/v) per membrane fraction (corresponding to cells cultured in one well of a six-well plate) for 15 min on ice. Subsequently, the solubilization mixture was centrifuged for 10 min at 4°C and 21,000 xg, and the supernatant was mixed with 90 μl of acquisition buffer containing 0.11% GDN (w/v). The diluted sample was then passed through a Zeba Spin desalting column (0.5 mM, 7 kDa cutoff; Thermo Fisher Scientific, France) equilibrated in acquisition buffer containing 0.005% LMNG (w/v), 0.0005% CHS Tris (w/v) and 0.005% GDN (w/v).

For smFRET experiments, HEK293T cells were cultured in T-75 surface: Cell+, 2-position screw cap flasks (Sarstedt). After treatment with a 28.5 μg/μl solution of poly-l-ornithine-hydrobromide in DPBS at 37°C for 15 min, wells were rinsed once with DPBS and a total of 10,000,000 cells were seeded in 10 ml of DMEM per flask 17 hours before transfection. Transfections were performed using 10 μg of total DNA per flask with a 1:1:1:1 ratio of vectors coding for mGlu2-A258TAA-C2KKXX, FLAG-SNAP-mGlu2-A248TAG-C1KKXX, pIRE4-PGK-ePrFRS and pNEU-hMbPylRS(AF)-4xU6M15 in 500 μl of JetPrime buffer. A total of 20 μl of JetPrime Reagent per flask were added and incubated for 25 min at RT. Last, the transfection mixtures were added to the cells and incubated at 37°C for 6 hours. After 6h, medium was replaced with 10 ml fresh DMEM supplemented with 0.2 mM TCOK and 0.3 mM PrF. The medium was further exchanged with fresh TCOK and PrF-enriched medium after 24 and 48 h. 72h after transfection SNAP-tag labelling was performed on adherent cells at 37°C and 5% CO_2_ for 2h using a final concentration of 300 nM SNAP-surface 488, excess dye was removed by three cycles of washing with DPBS without Ca^2+^ and Mg^2+^ at RT. TCOK labelling was performed right after SNAP-tag labelling using a final concentration of 1 mM Cy3B tetrazine in acquisition buffer (20 mM Tris-HCl pH 7.4, 118 mM NaCl, 1.2 mM KH_2_PO_4_, 1.2 mM MgSO_4_, 4.7 mM KCl and 1.8 mM CaCl_2_) incubated for 15 min at 37°C. Excess dye was removed by three cycles of washing with DPBS without Ca^2+^ and Mg^2+^ at RT. PrF labelling was performed last using a final concentration of 10 μM AF647-Picolyl-Azide, 1.5 mM Aminoguanidine, 1.98 mM BTTAA, 0.36 mM CuS0_4_, 2 mM Na-Ascorbate in acquisition buffer incubated for 15 min at 37°C. Excess dye was removed by three cycles of washing with DPBS without Ca^2+^ and Mg^2+^ at RT. Crude membrane fractions were prepared as describe above.

Receptors were solubilized using 100 μl of acquisition buffer containing 1% LMNG (w/v) and 0.1% CHS Tris (w/v) per membrane fraction (corresponding to cells cultured in one T-75 flask) for 30 min on ice. Subsequently, the solubilization mixture was centrifuged for 20 min at 4°C and 21,000 g, and the supernatant was mixed with 900 μl of acquisition buffer containing 0.11% GDN (w/v). The diluted sample was then loaded to a 200 μl anti-FLAG resin gravity column equilibrated with acquisition buffer and incubated at 4°C for 1h, then reapplied 2 times. The column was washed with 1ml acquisition buffer containing 0.01% LMNG (w/v), 0.001% CHS Tris (w/v) and 0.01% GDN (w/v), then 1ml acquisition buffer containing 0.005% LMNG (w/v), 0.0005% CHS Tris (w/v) and 0.005% GDN (w/v) and finally eluted with three times 120 μl acquisition buffer containing 0.005% LMNG (w/v), 0.0005% CHS Tris (w/v), 0.005% GDN (w/v) and 0.2 mg/ml FLAG peptide (Sigma). For acquisition, sample was diluted in acquisition buffer to a final of 0.0025% LMNG (w/v), 0.00025% CHS Tris (w/v), 0.0025% GDN (w/v) and 10 mM glutamate for the glutamate condition.

### Single molecule FRET acquisition

2-color smFRET experiments with pulsed interleaved excitation (PIE)–multiparameter fluorescence detection (MFD) were performed on a homebuilt confocal microscope using the SPCM 9.85 software (B&H) as described previously (Olofsson and Margeat 2013). Modifications are described in the following. A combination of 530/20 (530AF20, Omega Optical, Brattleboro, VT, USA) and 530/10 nm (FLH532-10, Thorlabs, Maisons-Laffitte, France) bandpass filters was used for Cy3B excitation. A 488/10 (Z488/10 X, Chroma, Bellows Falls, VT, USA) bandpass filters was used for SNAP-surface488 excitation. A 635/10 (FLH635-10, Thorlabs, Maisons-Laffitte, France) bandpass filter was used for AF647 excitation. Inside the microscope, the light was reflected by dichroic mirrors that match the excitation/emission wavelengths of the respective fluorophore combinations (Cy3B/AF647: FF545/650-Di01, Semrock, Rochester, NY, USA and SNAP-surface488/AF647: FF500/646-Di01, Semrock, Roch-ester, NY, USA) and coupled into a 100×, numerical aperture 1.4 objective (Nikon, France). The following emission filters were used: Cy3B parallel and perpendicular ET BP 585/65 (Chroma, Bellows Falls, VT, USA); AF647 parallel and perpendicular FF01-698/70-25 (Semrock, Rochester, NY, USA); AF488 parallel 535/50 BrightLine HC, perpendicular 530/43 BrightLine HC (Semrock, Rochester, NY, USA). Dual color emission was separated using FF649LP long pass filters (parallel and perpendicular, Semrock, Rochester, NY, USA) for Cy3B with AF647 and AT608LP (parallel, Chroma, Bellows Falls, VT, USA) together with FF560LP (perpendicular, Semrock, Rochester, NY, USA) for SNAP-surface488 with AF647.

3-color smFRET measurements were performed on a home-built confocal setup with pulsed-interleaved excitation (PIE) and multiparameter fluorescence detection (MFD) as described previously (Barth et al. 2019). Briefly, the triple labelled mGlu samples were diluted to 10-20 pM and measured for 1-3 hours in solution. The laser powers used during the experiments were 40μ for blue (485 nm), 30 μW for green (565 nm) and 15 μW for red (647 nm) lasers. The collected data were analyzed with the open source software package PIE analysis with MATLAB (PAM) (Schrimpf et al. 2018). Bursts were selected as described previously using a sliding time window approach on the total signal, requiring at least 8 photons per time window of 500 μs and at least 40 photons in total per burst (Eyal Nir, Xavier Michalet, Kambiz M. Hamadani, Ted A. Laurence, Daniel Neuhauser, Yevgeniy Kovchegov 2006). Triple-labeled bursts were selected based on stoichiometry thresholds (S_BG_, S_BR_, and S_GR_) using a lower boundary of 0.3 and an upper boundary of 0.8. To additionally remove photobleaching and blinking events, the ALEX-2CDE filter(Tomov et al. 2012) was applied, which was calculated pairwise for the three excitation channels. The 3-color PDA analysis was done as described before(Antonik et al. 2006; Barth et al. 2019). Detected fluorescence intensities from all three fluorophores were sued to calculate the uncorrected FRET efficiency (proximity ratios, PR) values. The underlying population for each PR histograms, a maximum likelihood estimator based on a three component Gaussian mixture is used. First, the PR_GR_ histogram is fitted with binomial distribution function. Then, the two- and three-dimensional description of the 3-color FRET data is done by binomial and trinomial distributions. For the apo state, a single state (R_o_) was used to described both the PR_GR_ and PR_BR_ histograms. In the presence of glutamine, both PR_GR_ and PR_BR_ histograms were fitted with contributions from 2 states (A_c_ and I_Glu_).

## Author contributions

LB: Designed mutagenesis. Performed mutagenesis, protein expression, labeling and purification, 2-color and 3-color single molecule FRET experiments. Analyzed and interpreted data. Wrote the manuscript draft.

EB: Performed 3-color smFRET measurements. Analyzed and interpreted data. CC: Designed and performed mutagenesis.

JPP: Designed the research strategy and acquired funds. Provided reagents. Interpreted data. PR: Designed the research strategy and acquired funds. Provided reagents. Interpreted data.

EM: Designed the research strategy. Acquired funds. Built the microscope and supervised the 2-color experiments. Interpreted data. Wrote the manuscript draft.

DCL: corresponding: Built the microscope and supervised the 3-colors experiments. Interpreted data. Edited the manuscript.

RBQ, corresponding: Designed the research strategy. Acquired funds. Designed ncAA incorporation strategies. Performed mutagenesis. Established protein labeling and solubilization. Analyzed and interpreted data. Wrote the manuscript draft.

## Acknowledgements

We thank the members of the IBM team (CBS, Montpellier) for fruitful discussions, the Arpège platform (IGF, Montpellier) for providing facilities and technical support, PerkinElmer Cisbio for providing reagents and the France-BioImaging national infrastructure. We further thank I. Coin, and R. Serfling (Universität Leipzig, Germany) for providing plasmids for ncAA incorporation. This work was supported by the INSERM recruitement dotation (RBQ), the “Agence Nationale pour la Recherche” (ANR 18-CE11-0004-02, EM and JPP) and the CBS2 doctoral school scholarship (LB). DCL gratefully acknowledges funding from the German Research Foundation (DFG) via the Sonderforschungsbereich 1035 (Projekt number 201302640, project A11 to DCL.) and the support from the Federal Ministry of Education and Research (BMBF) and the Free State of Bavaria under the Excellence Strategy of the Federal Government and the Länder through the ONE MUNICH Project Munich Multiscale Biofabrication. DCL also thankfully acknowledges the support of the Ludwigs-Maximillians-Universität München through the Center for NanoScience (CeNS) and LMUinnovativ BioImaging Network (BIN).

## Competing interests

The authors declare that they have no competing interests.

## Notes

### Competing Interest Statement

The authors have declared no competing interest.

