## Supplementary figures for "Triple labeling resolves a GPCR intermediate state by 3-color single molecule FRET"

[*]Corresponding authors

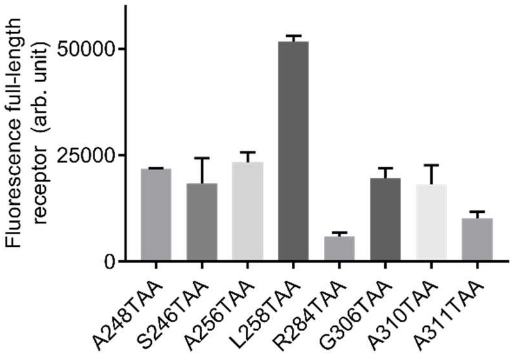


**Supplementary figure 1. Incorporation efficiency of TCOK in response to TAA suppression at different positions within the lower lobe of the VFT domain.** Incorporation was evaluation based total fluorescence measured from HEK293T cells expressing the indicated stop codon mutants in the presence of PylRS-$tRNA_{UUA}$ and TCOK. Labeling of full-length receptors, presented at the cell surface as a result of successful incorporation of TCOK, was achieved through N-terminal SNAP-tag labeling using the cell-impermeable Lumi4-Tb-SNAP substrate. Data are shown as the mean +/- SD from three individual measurements.

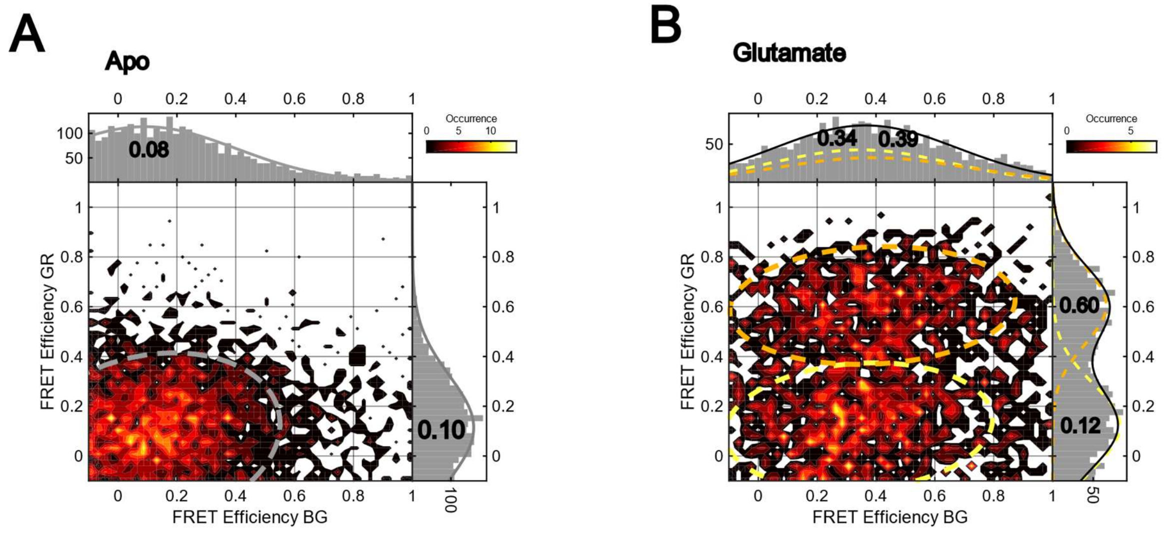


**Supplementary figure 2. 2-dimensional FRET efficiency plot of lower lobe (FRET efficiency GR) and diagonal (FRET efficiency BG) VFT domain sensors in the absence (A) and presence of glutamate (B) from 3-color smFRET acquisiton.** The FRET efficiency BG reports on the distance between the N-terminal SNAP-tag in one protomer and position 258 of the lower-lobe of the other protomer.
